# The anxiety and ethanol intake controlling GAL5.1 enhancer is epigenetically modulated by, and controls preference for, high fat diet

**DOI:** 10.1101/2020.07.08.186643

**Authors:** Andrew McEwan, Johanna Celene Erickson, Connor Davidson, Jenny Heijkoop, Yvonne Turnbull, Mirela Delibegovic, Chris Murgatroyd, Alasdair MacKenzie

**Affiliations:** School of Medicine, Medical Sciences and Nutrition, Institute of Medical Sciences, Foresterhill, University of Aberdeen, Aberdeen, Scotland, UK, AB25 2ZD; Manchester Metropolitan University, Manchester UK

**Keywords:** GAL5.1 enhancer sequence, CRISPR genome editing, galanin, EGR1, chromatin immunoprecipitation, gene regulation, 5mC/5hmC content, early life stress, fat preference, anxiety and substance abuse

## Abstract

Excess maternal fat intake and obesity increase offspring susceptibility to conditions such as chronic anxiety and substance abuse. We hypothesised that these susceptibilities are established through environmentally modulated DNA-methylation (5mC/5hmC) changes in regions of the genome that modulate mood and consumptive behaviours. We explored the effects of environmental factors on 5mC/5hmC levels within the GAL5.1 enhancer that controls anxiety-related behaviours and alcohol intake. We first observed that 5mC/5hmC levels within the GAL5.1 enhancer differed significantly in different parts of the brain. Moreover, we noted that early life stress had no significant effect of 5mC/5hmC levels within GAL5.1. By contrast, we identified that allowing access of pregnant mothers to high-fat diet (>60% calories from fat) had a significant effect on 5mC/5hmC levels within GAL5.1 in hypothalamus and amygdala of resulting male offspring. Cell transfection based studies using GAL5.1 reporter plasmids showed that 5mC has a significant repressive effect on GAL5.1 activity and its response to known stimuli such as EGR1 expression and PKC agonism. Intriguingly, CRISPR driven disruption of GAL5.1 from the mouse genome, although having negligible effects on metabolism, significantly decreased intake of high fat diet suggesting that GAL5.1, in addition to being epigenetically modulated by high-fat diet, also actively contributes to the consumption of high-fat diet suggesting its involvement in an environmentally-influenced regulatory-loop. Furthermore, considering that GAL5.1 also controls alcohol preference and anxiety these studies may provide a first glimpse into an epigenetically controlled mechanism that links maternal high-fat diet with transgenerational susceptibility to alcohol abuse and anxiety.

## Background

In addition to their involvement in obesity (1,2) there is also evidence that diets high in fat are associated with an increase in anxiety and substance abuse in subsequent human generations (3,4). These human studies are supported by observations that high-fat diet and maternal obesity increase substance abuse in the offspring of rats (5) and anxiety related behaviour in C57BL/6 mouse models (6). Therefore, in addition to the mechanisms regulating the intake of high-fat diets, there is also a priority to uncover the genomic and epigenetic mechanisms that link high-fat diet to anxiety and substance abuse disorders.

Nutrient intake (fats and ethanol) and mood are controlled by the expression of neuropeptides in regions of the brain that include the hypothalamus and amygdala (7). The expression of one of these neuropeptides; galanin, a 30 amino acid peptide encoded by the *GAL* gene, contributes to the regulation of fat intake in animals (2,8-10) plays a role in modulating mood (11-14) and regulates ethanol intake (15-18). Considering its important role in these processes, much remains to be determined regarding the mechanisms that regulate the tissue-specific expression of the GAL gene. In order to address this knowledge gap we previously used comparative genomics to identify a highly conserved enhancer sequence (GAL5.1) that lay 42 kilobases (kb) 5’ of the *GAL* gene transcriptional start site in humans and demonstrated its activity in galanin expressing cells of the hypothalamus including the PVN, dorsomedial hypothalamus (DMH) and arcuate nucleus (19,20). Analysis of two polymorphisms in the GAL5.1 enhancer using the UK Biobank suggested a mechanistic link between allelic variants of these polymorphisms and alcohol abuse when stratified against sex (male) and anxiety (20). Subsequent use of CRISPR genome editing, to disrupt the mouse GAL5.1 enhancer (mGAL5.1KO), demonstrated a role for mGAL5.1 in the tissue specific expression of the *Gal* gene in hypothalamus and amygdala as well as the modulation of alcohol intake and male anxiety-like behaviour (20).

Because of the known link between high-fat diet and increased susceptibility to anxiety and alcohol abuse in offspring of mothers fed high fat diet, we explored the hypothesis that the anxiety and alcohol intake modulating GAL5.1 enhancer could be epigenetically influenced by environmental stimuli that include early life stress and high-fat diet. We also explored the effects of DNA-methylation on GAL5.1 activity and its interactions with, and response to, known stimuli such as PKC agonism and EGR1 expression. Finally, we asked whether deleting the GAL5.1 enhancer has any effect on metabolism or fat intake. We discuss our findings in the wider context of a role for GAL5.1 genetics and epigenetics in modulating fat intake as well as the possible role of altered GAL5.1 activity in increasing susceptibility to anxiety and alcohol abuse in future generations.

## Methods

### Animal studies

All animal studies were performed in full accordance with UK Home Office guidelines. Male and female homozygous wildtype and mGAL5.1KO age matched littermates were single housed under standard laboratory conditions (12 h light/12 h dark cycle), in plastic cages with food and water available ad libitum, depending on the experiment.

### Epigenetic effects of early life stress by maternal deprivation

Wildtype C57BL/6 females were housed with wild type C57BL/6 males and allowed ad-libitum access to standard CHOW diet. Once females became pregnant males were removed and females were allowed to litter down. Immediately after birth mothers were removed from the cages for two hours a day for the first 12 days as previously described (21). Once weaned litters were humanely sacrificed by euthatol injection and brain tissues (hypothalamus, hippocampus and amygdala) recovered and rapidly frozen on dry ice.

### Epigenetic effects high-fat diet studies

Wildtype C57BL/6 wild type animals were housed with wild type C57BL/6 males and allowed ad libitum access to a choice of low fat diet (LFD; 22.03 kcal% protein, 68.9kcal% carbohydrate and 9.08 kcal% fat) or high fat diet (HFD; 20 kcal% protein, 20 kcal% carbohydrate and 60 kcal % fat; Research Diets Inc.) in different hoppers. Both hoppers were weighed regularly to ensure intake of HFD and LFD. Once females became pregnant males were removed and females allowed to litter down. After weaning (3-4weeks), animals were humanely sacrificed by euthatol injection and brain tissues (hypothalamus, hippocampus and amygdala) recovered and rapidly frozen on dry ice.

### DNA extraction bisulfite conversion and pyrosequencing

Genomic DNA was extracted and purified with the AllPrep DNA/RNA Mini Kit (Qiagen) and optimized for 10-15 mg brain tissue as per manufacturer’s instructions. The concentration and purity of genomic DNA was determined using a Nanodrop One^C^ (Thermo) and 500 ng of genomic DNA was bisulphite-converted using the EpiMark Bisulfite Conversion Kit (New England BioLabs) as per manufacturer’s instructions. The mouse GAL5.1 enhancer region analysed (chr19:3,441,054-3,441,445, GRCm38/mm10) contained 8 CpG sites (see Figure 1). Primers were used to amplify two regions of the mouse Gal5.1 enhancer covering 8 CpGs: CpGs 4-8 (F– TTTAGTAGAGGAAATAAAATAGTAGAAAAA-Biotinylated; R: CCCCAAAAAACCACAAAACCTA) CpGs 9-11 (F – GGATGGAGGAATTTTTTTGTGTT; R: CCCCAAAAAACCACAAAACCTA -Biotinylated) using MyTaq HS mix PCR reagents (Bioline). Amplicons were processed on the Qiagen Q24 Workstation and sequenced in duplicate on the Qiagen Q24 pyrosequencer using the sequencing primers CpGs 4-8 (AACAATTTAAACAAAAAATAACATT) CpGs 9-11 (GAGTTTGATTGATAATAGTAGTAT) and DNA methylation levels across each CpG measured. These were expressed as percentage methylation.

**Figure 1.**
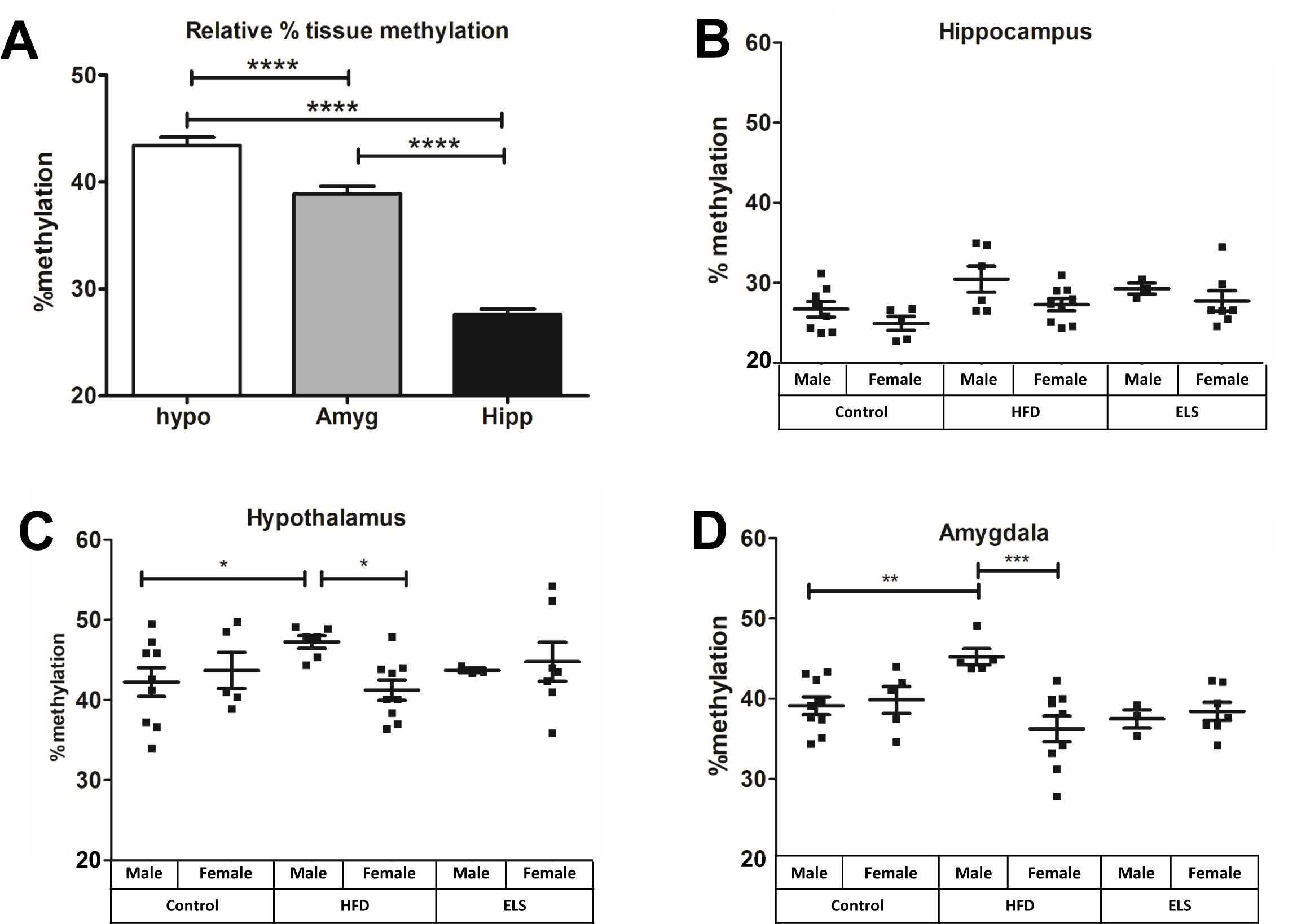
Levels of DNA-methylation (5mC) within GAL5.1 differs significantly between specific tissues and is affected by maternal access to high fat diet but not early life stress. **A**. bar plot demonstrating differential levels of 5mC that occur within the GAL5.1 enhancer in DNA derived from hypothalamus (hypo), amygdala (amgd) and hippocampus (hipp). **B-D**, scatterplots showing the effects of high-fat diet (HFD) and early life stress (ELS) on 5mC levels of GAL5.1 derived from hippocampus (**B**), hypothalamus (**C**) and amygdala (**D**). n.s.; not significant, *; p<0.05, **; p<0.01, ***;p<0.005.

### Chromatin Immunoprecipitation analysis

SH-SY5Y cells grown to confluence in T-25 flasks were transfected with 0.25pM of each reporter construct (pLucGAL5.1GG or pLucGAL5.1ΔECR1(20)) or an expression construct (empty pcDNA3.1 vector or pcDNA-*Egr1*) using the JetPrime™ Transfection Reagent according to manufacturer instructions (Polyplus-Transfection, France). PKC activity was stimulated in these cell cultures using Phorbol 12-Myristate 13-Acetate (PMA) to a final concentration of 100nM. After 24 hours cells were then fixed with a 1% (v/v) formaldehyde solution supplemented with cOmplete™ Mini EDTA-free Protease Inhibitor Cocktail (Roche®, UK) for 15 minutes with gentle agitation at room temperature. Chromatin immunoprecipitation analysis was then undertaken as previously described (22) using a ChIP-Certified EGR1 Antibody (<u>MA5-15008, ThermoFisher Scientific®,</u> <u>UK</u>) and an isotype-matched Normal Rabbit IgG (<u>12-370, Sigma-Aldrich®, UK</u>) as negative control. Immunoprecipitated DNA was analysed for relative enrichment of EGR1 bound GAL5.1 using qPCR within a LightCycler® 480 (Roche®, UK) using forward and reverse primers specific for the human GAL5.1 enhancer (hGal5.1_Plasmid_F; 5’-GCTAAAGCGGACTAGGGTCATTGTAAG-3’, hGal5.1_Plasmid_R; 5’-GAACACGCAGATGCAGTCGG-3’). In order to control for the amount of pLucGAL5.1GG, and pLucGAL5.1ΔECR1 plasmid transfected into each cell culture parallel qPCR reactions were undertaken using primers to detect the luciferase gene (hLuc_Plasmid_F; 5’-GCCGTTGTTGTTTTGGAGCA-3’, hLuc-Plasmid_R; 5’-GCGGTTGTTACTTGACTGGC-3’) from pre-immunoprecipitated cell lysates as a transfection control to normalise the qPCR data derived from the ChIP assay.

### Reporter assays

The human GAL5.1 enhancer was cloned into the pCpGfree-promoter-lucia vector (InvivoGen; contains the EF1α promoter) and subjected to CpG methylation using SssI (New England Biolabs) as described in manufacturer’s instructions. Methylated and unmethylated plasmids were transfected into SHSY-5Y cells vector using Jetprime transfection reagent (Polyplus-transfection) and incubated for 24 hours. Cells were lysed using passive lysis buffer and the activity of the enhancer assessed using a Lucia QUANTI-Luc Gold Assay #3 luciferase kit (InvivoGen). Because of clear evidence of the response of the pGL4.74 Renilla normalisation control plasmid to PMA treatments and co-transfection with the EGR1 expression vector Lucia activity levels were normalised against quantitative PCR (qPCR) of levels of Lucia gene DNA present in cell extracts recovered using Qiagen PCR cleanup kits (Qiagen). Lucia luciferase activity was normalised against the crossing point (Cp) values generated from quantitative polymerase chain reaction (qPCR) of the lucia reporter vector. Absolute quantification of the transfected lucia reporter vector was determined in order to verify transfection efficiency between all samples. After media from cell wells was extracted, the cells were lysed using 100µl passive lysis buffer (Promega) per well. Total DNA was isolated from the cell lysates using the QIAquick® PCR Purification Kit (QIAGEN). The following primers used were designed to the lucia DNA sequence for amplification of the reporter gene (JE041; 5’-GAAATCAAGGTGCTGTTTGC-3’, JE042; 5’-TATCATCTGTCCCCAGCC-3’). Samples were loaded into an opaque 384-well plate in triplicate following the standard cycling parameters of a SYBR Green protocol with 60°C annealing temperature. Amplification and quantitative analysis were carried out using the LightCycler® 480 System (Roche).

### High fat preference studies

Mice containing a disrupted GAL5.1 enhancer (20) were maintained as a colony on a heterozygous C57BL/6 background which was mated to produce homozygous wild type and homozygous GAL5.1 KO age matched and sex matched individuals. These were identified using PCR of earclip DNA (20). Once identified, animals were assigned random numbers to hide their genotype from the operators of subsequent tests. For high fat diet preference studies, singly housed animals were provided with a choice of LFD or HFD in different hoppers. The position of each hopper was changed regularly to rule out the possibility of position effects. Animals were weighed daily over a period of 23 days and LFD and HFD were also weighed daily to determine intake of each diet.

### Metabolic analysis

Wild type and GAL5.1KO animals were maintained for one week in sealed TSE metabolic cages where calorie intake (kcal/hour), O_2_ consumption (ml/hr), energy expenditure (kcal/hour) locomotor activity, CO_2_ production (ml/hour), energy balance (kcal/hr) respiratory exchange ratios and locomoter activity (beam breaks) were continuously monitored for 144 hours with readings taken at 30 minute intervals.

### Data analysis

From *in-vivo* pilot studies we calculated that a minimum of 6-8 animals per group would enable detection of a 25% difference between different parameters (e.g. High Fat Diet, weight gain, metabolism) with 80% power using one-way ANOVA and/or general linear modelling. Statistical significance of data sets were analysed using either two way analysis of variance (ANOVA) analysis with Bonferroni post hoc tests or using one tailed or two tailed unpaired parametric Student *t*-test as indicated using GraphPad PRISM version 5.02 (GraphPad Software, La Jolla, CA, USA).

## Results

### Levels of CpG methylation (5mC/5hmC) within GAL5.1 varies significantly between brain regions

We have previously shown that the GAL5.1 enhancer controls anxiety and alcohol intake (20). This provided us with a unique opportunity to determine the effects of environmental factors such as dietary changes and stress on 5mC/5hmC levels within an enhancer region shown to control behaviours with a direct effect on human health. We first asked whether levels of DNA-methylation (5mC/5hmC) within GAL5.1 differed between brain regions or were similar. To address this question, DNA from 3 different brain tissues (hypothalamus, amygdala and hippocampus) was recovered from mouse pups and bisulfite converted to detect levels 5mC and 5hmC within GAL5.1 analysed using pyrosequencing. We observed that 5mC/5hmC levels within GAL5.1 varied considerably between different brain tissues such that in hippocampus 5mC levels only reached 23-32% within GAL5.1. However, in amygdala and hypothalamus 5mC levels were between 35 and 50% (**Figure 1A**).

### Levels of CpG methylation (5mC/5hmC) within GAL5.1 is not affected by early life stress

Because changes in CpG methylation (5mC/5hmC) is known to occur within enhancers as a result of early life stress and diet (21,23), we subjected newborn wildtype C57BL/6 mouse pups to maternal deprivation for 2 hours a day for the first 12 days of their lives as previously described (21). After weaning of these mice, DNA from 3 different brain tissues (hypothalamus, amygdala and hippocampus) was recovered, bisulfite converted and levels of 5mC/5hmC within GAL5.1 analysed using pyrosequencing. We did not observe any significant changes in 5mC/5hmC levels in any of the tissues recovered from animals subjected to early life stress (**Fig 1B-D**).

### GAL5.1 displays significantly altered 5mC/5hmC levels in animals whose mothers had access to HFD

We provided pregnant mice with a choice of low-fat or high-fat diet during pregnancy and during the rearing of their pups. Following weaning, pups were humanely sacrificed and genomic DNA was recovered from tissues dissected from hypothalamus, hippocampus and amygdala. This DNA was subjected to bisulfite conversion and pyrosequenced to determine changes in 5mC levels within the GAL5.1 enhancer in the presence of access to HFD. We were unable to detect any changes in GAL5.1 5mC levels in DNA derived from hippocampal tissues of animals whose mothers had access to high-fat diet (**Fig 1B**). However, significant changes in CpG methylation were observed in DNA derived from the hypothalamic regions of animals whose mothers had been exposed to high-fat diet such that there was a significant increase in methylation in males and a clear divergence in methylation between males and females (**Figure 1C**). This increase in male GAL5.1 5mC levels and sexual divergence was more pronounced in DNA derived from amygdala (**Figure 1D**).

### The EGR1 transcription factor physically interacts with the GAL5.1 enhancer at a highly conserved EGR1 binding site

We have previously shown that the GAL5.1 enhancer responds to expression of the early growth response factor (EGR1) transcription factor protein (20). However, these experiments stopped short of establishing a direct physical interaction. In order to determine their relationship we transfected SHSY-5Y neuroblastoma cells with equimolar equivalents of a luciferase vector containing either the complete human GAL5.1GG sequence (pLucGAL5.1GG) or GAL5.1GG lacking 10 base pairs corresponding to the highly conserved EGR1 binding site within GAL5.1(pLucGAL5.1ΔEGR1). It is important to point out that the aim of this experiment was not to determine luciferase activity but to determine levels of EGR1 interaction with GAL5.1 using the luciferase gene as a transfection control. In parallel, we also transfected an empty expression vector (pcDNA) or the same vector also expressing the cDNA of the EGR1 transcription factor (pcDNA-EGR1). We also treated these cells with the PKC agonist PMA. After incubation for 24 hours we used a ChIP-verified EGR1 antibody on cell lysates derived from transfected SHSY-5Y cells to immunoprecipitate DNA which had bound to the EGR1 protein and, after reversing cross links, assayed the amounts of GAL5.1 DNA that had formed a complex with EGR1. Using primers specific for the human GAL5.1 enhancer we used qPCR to demonstrate that the anti-EGR1 antibody immunoprecipitated significant amounts of GAL5.1 enhancer DNA and that this interaction increased in the presence of EGR1 expression or PKC activation (**Fig 2A**). In contrast, we were unable to immunoprecipitate little, if any, GAL5.1 DNA from cells which had been transfected with the plasmid missing the 10bp containing the highly conserved EGR1 binding site (**Fig 2B**). Transfections were normalised using quantitative PCR against the firefly luciferase gene. We concluded that EGR1 physically interacted with GAL5.1 and that this interaction was stimulated by PKC signalling. In addition, our data also suggests that EGR1 binding within GAL5.1 only occurred within the highly conserved EGR1 binding site.

**Figure 2.**
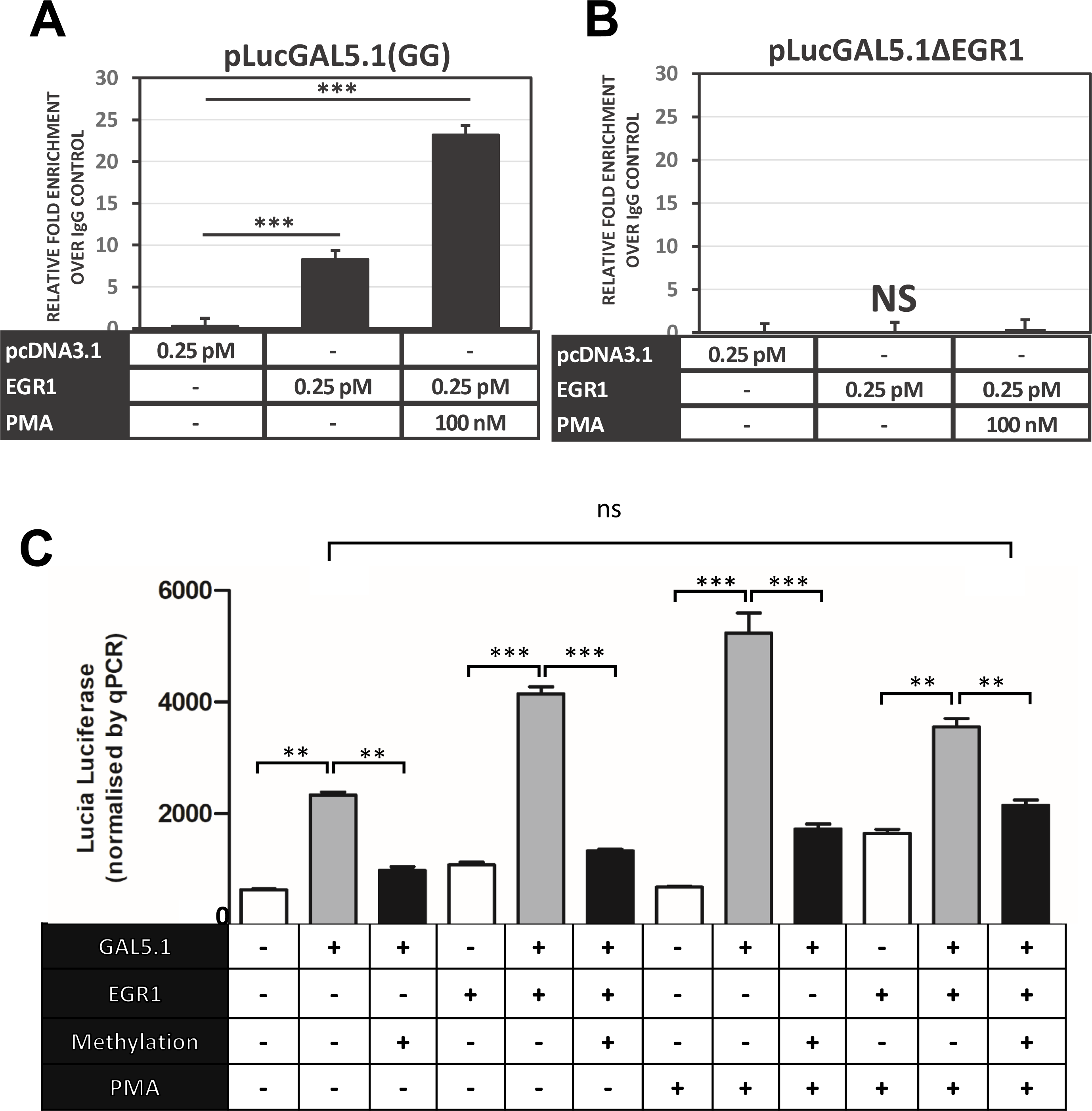
PKC agonism and DNA methylation modulate site specific interactions of the EGR1 transcription factor with GAL5.1. (**A and B**) Bar graphs showing enrichment of GAL5.1 DNA, from transfected SHSY-5Y cell lysates immunoprecipitated using a ChIP certified EGR1 antibody, as assessed by qPCR. SHSY-5Y cells were transiently transfected with either (**E**) the pLucGAL5.1GG plasmid or (**F**) the pLucGAL5.1ΔEGR1 combined with an empty expression vector (pcDNA3.1) or an expression vector expressing the EGR1 transcription factor protein (EGR1). Cultures were also treated with a PKC agonst (PMA). Data has been presented as the fold enrichment of immunoprecipitated GAL5.1 divided by the signal from the negative IgG control and normalised against luciferase gene DNA as a transfection control. n=3, error bars=SEM, ***; p<0.005, ns; not significant. **C.** A bar graph demonstrating relative Lucia Luciferase activity of the pCpG-free vector (white bars), the pCpGfree-GAL5.1 (Grey bars) and pCpGfree-GAL5.1 methylated using SssI (Black bars) and co-transfected with the pcDNA-EGR1 expression vector (EGR1; bars 3-5 and 10-12) or treated with the PKC agonist PMA (PMA; bars 7-12). Each Lucia luciferase values was normalised against the relative quantities of Lucia luciferase DNA detected in each cell extract using quantitative PCR. *n*=4, n.s.; not significant, **; p<0.01, ***;p<0.005.

### Methylation of the GAL5.1 enhancer suppresses its activity and its response to EGR1 binding and PKC activation

We next asked what effects CpG methylation would have on the activity of the GAL5.1 enhancer, its response to EGR1 expression and PKC activation which we have shown activates GAL5.1 (19,20). We first cloned the human GAL5.1 (GG) enhancer into the pCpGfree-promoter-Lucia luciferase vector that lacks CpG dinucleotides and contains the EF-1α promoter. We then subjected this vector to CpG methylation using the bacterial SssI enzyme and transfected methylated or unmethylated plasmid into SHSY-5Y neuroblastoma cells in the presence of an expression vector expressing the EGR1 transcription factor **(Figure 2C).** Levels of plasmid methylation were monitored using digestion by the HpaII enzyme which is sensitive to methylated CpG sites (**Figure S1**). These cells were then cultured for 24 hours in the absence or presence of the PKC agonist PMA. Due to evidence of trans-interactions between our renilla control and the pCpG-lucia vector (probably through the EF-1α promoter), we devised an alternative transfection control based on quantitative PCR of the Lucia reporter gene. We observed that the GAL5.1 enhancer increased expression of Lucia luciferase compared to the empty vector but that this increase was negated by CpG methylation (**Figure 2C**). The previously reported stimulatory effects of either EGR1 co-expression or activation of the PKC pathway (19,20) were similarly reduced by GAL5.1 methylation (**Figure 2C**). Interestingly, we observed that the activity of the non-methylated GAL5.1 enhancer was not significantly different from that of the methylated enhancer co-stimulated by both EGR1 expression and PKC activation (**Figure 2C**) suggesting that, together, EGR1 and PKC stimulation can overcome much the effects of 5mC on GAL5.1 activity.

### mGAL5.1KO mice do not demonstrate significant changes in weight gain or intake of CHOW diet

We had previously shown that CRISPR deletion of GAL5.1 from the mouse genome produced mice that drank significantly less ethanol and suffered less anxiety (19,20). In order to explore whether disruption of the GAL5.1 enhancer in mice had any significant effect on low fat food intake (CHOW) or weight gain we monitored the weight gain and CHOW intake of male and female homozygous wild type (WT) and GAL5.1KO animals. We found that, after 18-24 weeks, neither male nor female animals demonstrated significant differences in weight gain compared to wild type animals (**Figure 3A**). We also found that GAL5.1KO animals ate similar levels of CHOW diet to WT animals (**Figure 3B**).

**Figure 3.**
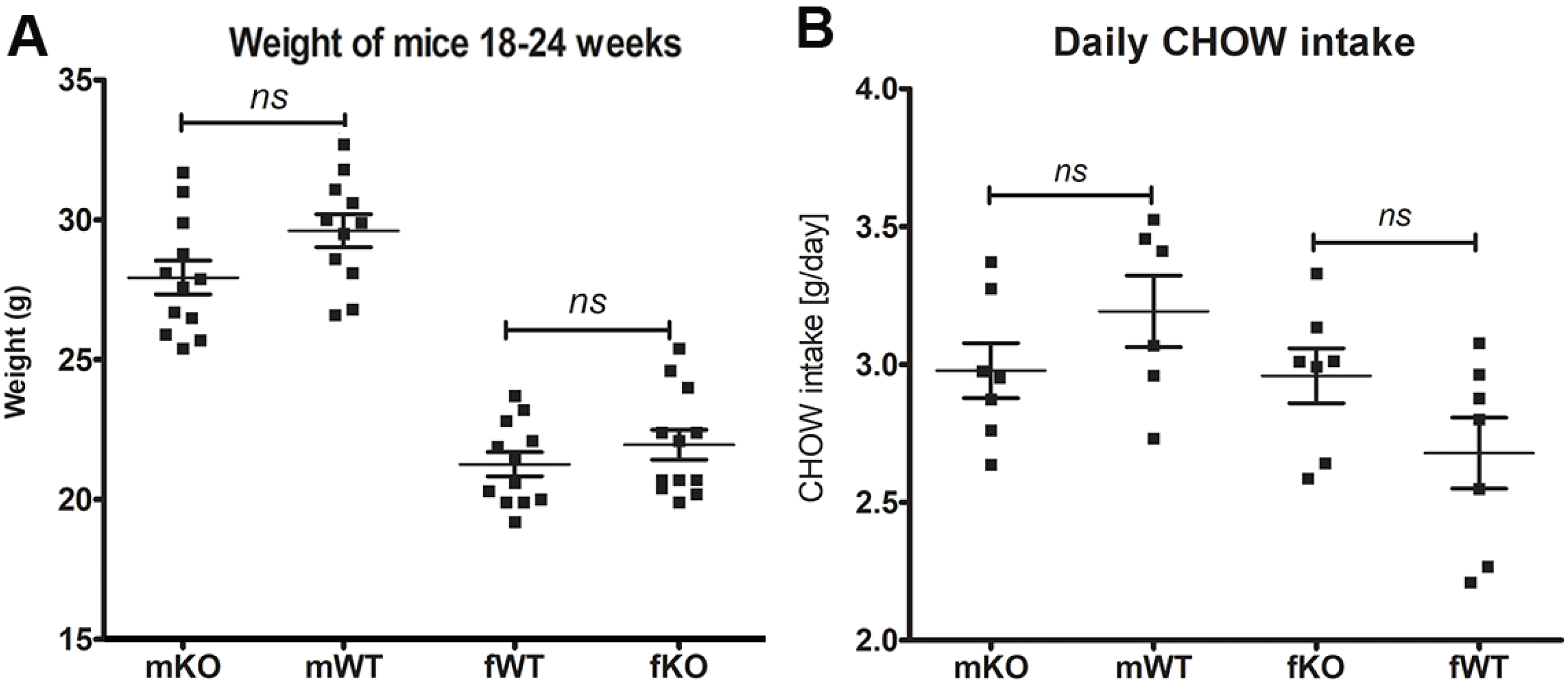
GAL5.1KO animals do not differ significantly from wild type animals in food intake or weight gain. **A**. Scatterplots showing a comparison of weights of male (m) and female (f) GAL5.1KO (KO) and wild type (WT) animals at 24 weeks of age. **B**. A comparison of daily CHOW intake of male and female wild type and knock out animals. ns=not significant.

### mGAL5.1KO mice do not demonstrate significant metabolic differences to WT animals

In order to explore the possibility that disruption of GAL5.1 significantly altered metabolism we subjected mGAL5.1KO and wild type male and female littermates to 116 hours of metabolic analysis using sealed TSE cages that monitored variables such as O2 consumption, CO2 production (ml/hour), energy expenditure ration (kcal/hour), energy balance (Kcal/hour). Over the 116 hours of the analysis we detected little or no significant difference in O_2_ consumption (**Fig 4A)**, CO_2_ production (**Fig 4B)** or energy expenditure (**Fig 4C**) in male and female mice. However, we detected a significant decrease in the respiratory exchange ratio (**Fig 4D**) in male animals that suggested higher activity levels. We observed an increase in distance travelled by male GAL5.1KO animals (**Figure 4E**) in addition to an increase in overall speed (**Figure 4F**) consistent with higher levels of exploratory behaviour associated with the reduced anxiety phenotype previously reported (20).

**Figure 4.**
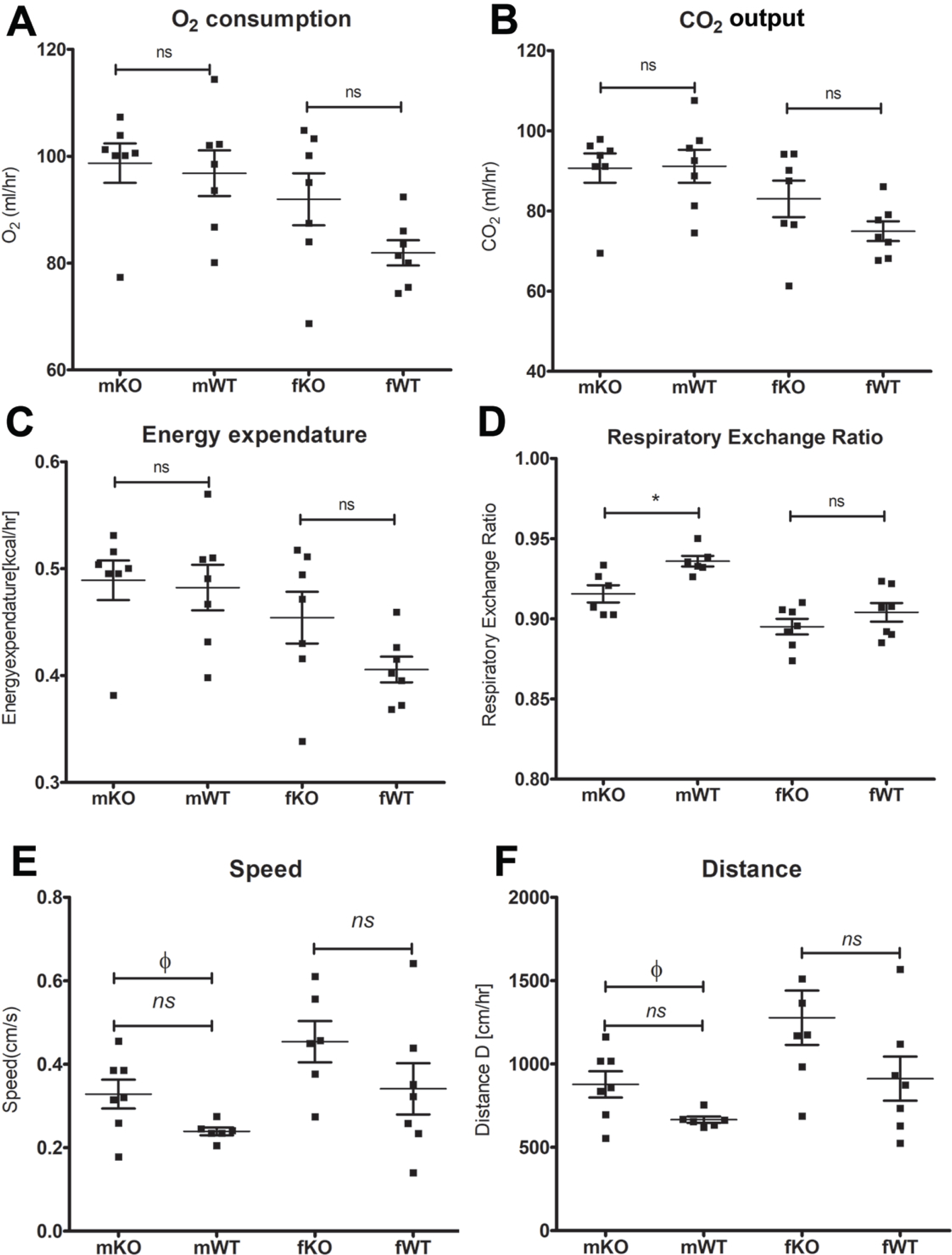
GAL5.1KO mice do not differ significantly in their metabolism. A-D; Scatterplots showing a comparison of average O2 consumtion (**A**), CO2 output (**B**), energy expenditure (C) and respiratory exchange ratio (D) in male (m) and female (f) GAL5.1KO (KO) and wild type (WT) animals recorded using sealed TSE cages. E, average speed (cm/second) and (F) distance travelled per hour (cm/hour). *; p<0.05, n.s. not significant (ANOVA), Φ= significant (2-tailed t-test).

### mGAL5.1KO mice exhibit decreased preference for high-fat diet

Previous studies demonstrated that deletion of exons 2-6 of the Gal gene in 129Ola/Hsd mice using ES-cell targeting (24) caused a significant reduction in the intake of high fat diet in these animals compared to wild type littermates but had no significant effects on protein or carbohydrate intake (10). Because we have previously shown that disruption of the GAL5.1 enhancer resulted in a significant decrease of Gal mRNA expression in hypothalamus we tested the hypothesis that CRISPR disruption of mGAL5.1 would affect intake of HFD in these mice. We provided singly housed, age- and sex-matched, littermate wild type and mGAL5.1KO animals with a choice of low fat diet (LFD; 6% of calories from fat) or high fat diet (HFD; 60% of calories from fat) and monitored intake of LFD and HFD over 23 days. Both male and female mGAL5.1KO mice consumed significantly less HFD overall compared to wild type littermates (**Fig 5A).** Analysis of the total intake of LFD demonstrated no significant differences between the intake of wild type and mGAL5.1KO animals (**Fig 5B**).

**Figure 5.**
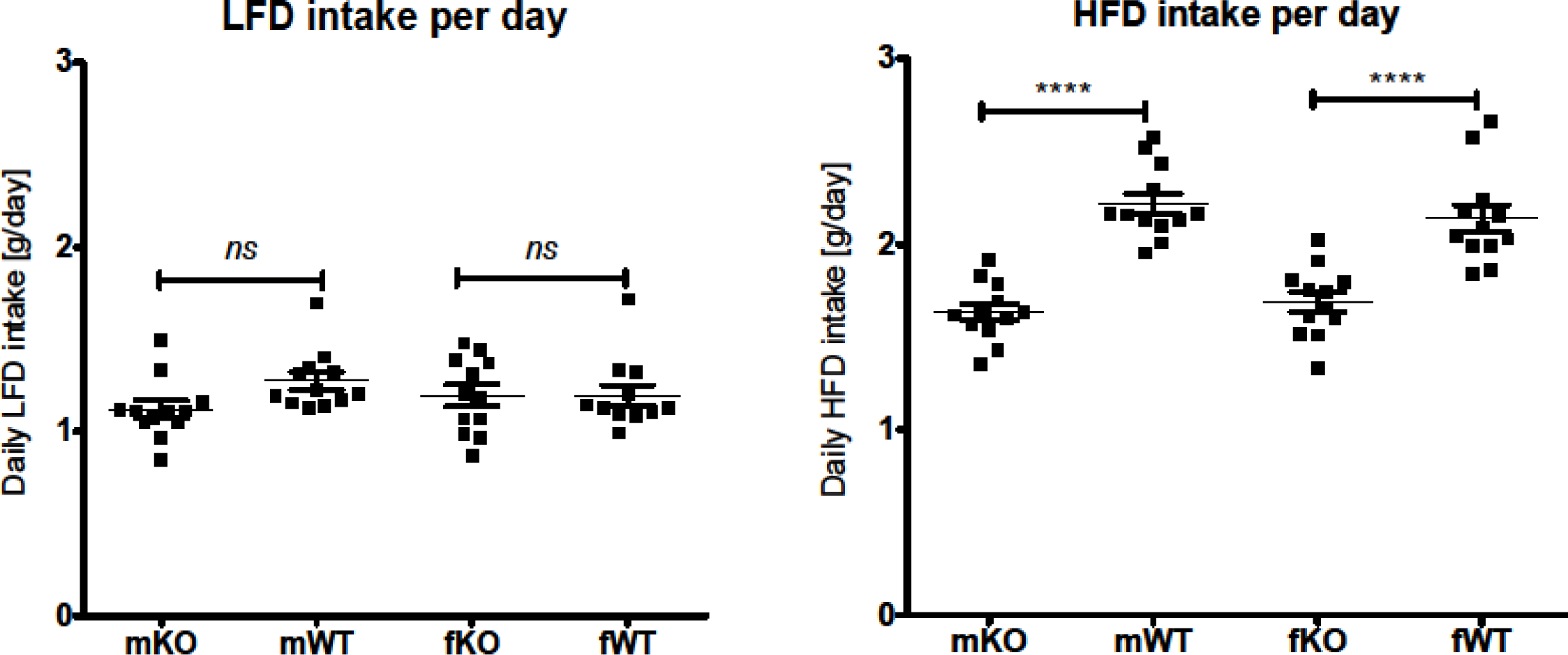
GAL5.1KO mice consume significantly less high-fat diet. A and B; scatterplots comparing daily intake of (A) high-fat diet and (B) low-fat diet comparing male (m) and female (f) GAL5.1KO (KO) and wild type (WT) mice. ns= not significant, ****; p<0.001.

## Discussion

Studies have suggested that increased maternal fat intake and obesity in both humans (3,4) and animal models (5,6) increase susceptibility to substance abuse and increased anxiety in resulting offspring. In the current study we explored the hypothesis that a contributing mechanism linking maternal high-fat diet and anxiety/substance abuse in offspring could involve the epigenetic modification of tissue specific enhancer regions through DNA-methylation to form 5-methylcytosine (5mC) or 5-hydroxy-methyl cytosine (5hmC) (25-27). Our decision to explore a role for 5mC/5hmC in this process was based on previous studies were environmental challenges such as high-fat diet and early life stress have been shown to change levels and distributions of 5mC/5hmC in the genomes of experimental animals (21,28) and alter enhancer activity through interaction with DNA binding proteins (29).

We focussed our attentions on the methylation of an enhancer sequence; called GAL5.1, that we have previously shown is responsible for supporting expression of the *GAL* gene in the hypothalamus and amygdala and which also controls ethanol intake and anxiety in males (19,20). We first asked whether environmental factors associated with poverty and deprivation in Western countries; namely early life stress by maternal deprivation or consumption of high fat diet, altered levels of DNA-methylation within the 11 CpG dinucleotides contained within the GAL5.1 enhancer sequence. We first noted significant differences in 5mC/5hmC levels in GAL5.1 derived from different tissues such that GAL5.1 from hippocampal tissues had the lowest 5mC/5hmC levels and hypothalamus and amygdala demonstrated the highest levels. These are interesting observations as the vast majority of large human cohort epigenome studies have based their analysis of levels and distribution of 5mC/5hmC on DNA derived from peripheral blood. However, given the observed differences in 5mC/5hmC levels observed between different brain tissues in the current study we suggest that caution should be exercised when interpreting studies based on the extrapolation of 5mC/5hmC levels in DNA derived peripheral blood to specific regions of the brain.

Although the repressive effects of 5mC on promoter activity through mechanisms involving the recruitment of methyl-CpG-binding proteins (MBD) and histone deacetylase (HDAC) containing complexes are well established and accepted (30,31) the same cannot be said for remote tissue specific enhancer regions. For example, using DNAse hypersensitivity; a widely accepted marker of active enhancers, the ENCODE consortium only observed a decrease in chromatin accessibility in 20% of accessible sites as a result of CpG methylation suggesting that the remaining 80% of DNAseI hypersensitivity sites were still active despite increased 5mC levels (32). Furthermore, an analysis of the effects of 5mC on the ability of 542 transcription factors (TFs) to bind DNA; an essential component of enhancer activation; found that only 23% of TFs experienced reduced binding as a result whereas the binding of 34% of TFs was actually increased by 5mC and the remaining TFs were unaffected (33). The role of DNA-methylation in influencing enhancer activity is further complicated by the influence of intermediate oxidation products of 5mC; that includes 5-hydroxymethyl-cytosine (5hmC), the product of the Ten-Eleven Translocation (TET) dioxygenase enzymes (34). 5hmC sites are attractive to transcription factors that activate transcription (35). 5hmC has been associated with active enhancers in stem cells (36,37) and deletion of TET2 from mouse stem cells reduces 5hmC levels and activity of enhancers (37). It is interesting, in this context, that the bisulfite sequencing protocol used in the current study is unable to differentiate 5mC from 5hmC so that it is as likely that the increased levels of 5mC observed within GAL5.1 as a consequence of maternal HFD intake actually reflects an increase in 5hmC levels. The possible role of 5hmC in upregulating enhancer activity, and our inability to differentiate 5mC and 5hmC using bisulfite conversion, may also explain the observations of increased 5mC/5hmC levels in hypothalamus and amygdala compared to the hippocampus that parallel the levels of Gal mRNA expression, which are significantly higher in hypothalamus and amygdala compared to hippocampus. Alternative methods of differentiating between genomic 5mC and 5hmC are now available which could help clarify the situation.

Previous analysis of the transcription factors that influence activity of GAL5.1 suggested the involvement of the EGR1 transcription factor by virtue of the fact that co-expression of EGR1 induced an up-regulation of transcriptional activity driven by the GAL5.1 enhancer (20). However, this evidence was circumstantial and could not rule out the possibility that the effects of EGR1 expression on GAL5.1 activity were indirect. We devised a unique experiment to determine whether the EGR1 protein directly interacted with the GAL5.1 enhancer at a predicted and highly conserved EGR1 binding site and whether this interaction was influenced by PKC activation. This involved transfecting neuroblastoma cells with DNA constructs containing the GAL5.1 enhancer containing, or lacking, the EGR1 binding site, in combination with a plasmid vector expressing the EGR1 protein. It is important to note that the luciferase gene in this experiment was only used as a normalisation control for transfection efficiency. We then carried out chromatin-immunoprecipitation analyses on nuclear lysates derived from these cell cultures using ChIP verified EGR1 antibodies. We found that, in the presence of the EGR1 expression vector, more GAL5.1 DNA was recovered strongly verifying a direct molecular interaction between EGR1 and GAL5.1. In contrast, deletion of the 10bp containing the EGR1 conserved consensus sequence greatly reduced recovery of GAL5.1 DNA demonstrating that this sequence within GAL5.1 was required for GAL5.1-EGR1 interaction. Removal of this 10bp EGR1 binding site also blunted the ability of PKC activation to induce EGR1 binding. These unique observation add to the findings of our previous studies (20) by demonstrating a direct interaction between the EGR1 transcription factor and a 10bp sequence within the GAL5.1 enhancer.

We next designed another unique experiment to determine the effects of 5mC on the activity of the GAL5.1 enhancer and its response to EGR1 binding and PKC agonism. To achieve this we cloned the GAL5.1 enhancer into a luciferase reporter vector (pCpGfree-lucia) that contained no CpG dinucleotides. This allowed us to selectively methylate CpG dinucleotides within the GAL5.1 enhancer, using the SssI enzyme, without affecting the vector backbone. These studies showed that CpG methylation of the GAL5.1 enhancer has a significant repressive effect on its activity in SHSY-5Y cells and confirmed previous studies demonstrating the repressive effects of 5mC on regulatory activity. However, our studies could not explore the effects of 5hmC on enhancer activity, which has been associated with increased enhancer activity (34-37). Thanks to the recent availability of recombinant TET-proteins, that are able to convert 5mC to 5hmC, it may be possible to compare the effects of 5mC and 5hmC on GAL5.1 action in the near future.

Previous studies have shown that the binding affinity of the EGR1 transcription factor to DNA is not affected by 5mC (38). However, we observed that 5mC suppressed GAL5.1 activity even in the presence of EGR1. This suggests that the binding of another transcription factor; who’s binding to DNA is affected by CpG 5mC, is critical to the normal functioning of GAL5.1. Identifying this transcription factor will be a major goal of subsequent analyses. 5mC CpGs within GAL5.1 also have a significant impact on the response of GAL5.1 to activation of the PKC pathway. Thus, the ability of 5mC to repress PKC activation of GAL5.1, even in the presence of EGR1, represents further evidence for the need of a second 5mC sensitive transcription factor in the normal response of GAL5.1. Interestingly, levels of activity of the fully methylated GAL5.1 enhancer in the presence of both EGR1 expression and PMA do not differ significantly from that of the unmethylated GAL5.1 enhancer in the absence of these stimuli suggesting that the effects of 5mC on GAL5.1 might be overcome by increased TF binding and stimulation of signal transduction pathways *in vivo.* In the context of the role of the EGR1 transcription factor in modulating levels of 5mC/5hmC at the GAL5.1 locus it is also interesting that recent studies have identified EGR1 as a major recruiter of TET proteins to specific loci (39) a consideration which we will also consider in the design of future studies.

In the wider context of the relationship of maternal high-fat diet to increased susceptibility to substance abuse and anxiety, the current study suggests the possible mechanistic involvement of the GAL5.1 enhancer in this process. We have previously shown that GAL5.1 governs ethanol intake and anxiety related behaviour in mice that is mirrored by a significant association between alcohol abuse and increased anxiety in humans. In the current study we show that high-fat diet causes a significant change in 5mC/5hmC levels that are known to affect enhancer activity and that, in turn, GAL5.1 governs the decision to eat high-fat diet but also affects anxiety and alcohol intake. Taken together, we propose that maternal high-fat diet induced changes in GAL5.1 5mC/5hmC levels may alter GAL5.1 activity in subsequent generations in such a way as to affect the ability of GAL5.1 not just to alter fat intake, but to also affect anxiety and the decision to drink alcohol.

## Conclusions

Whilst much remains to be done to differentiate the effects of environmental influence on the distribution of 5mC and 5hmC in GAL5.1 and how 5mC and 5hmC differentially affect GAL5.1 activity at a tissue specific level, the fact remains that we have identified a compelling epigenetic mechanism that may link high-fat diet to the modulation of anxiety and alcohol intake. These unique studies also provide an important stepping stone in our voyage to discover the influence of environmentally modulated regulatory mechanisms in the development of neuropsychiatric disorders

## Acknowledgements

A. R. M. was funded by BBSRC project grant (BB/N017544/1).

## References

1. Liu, D., Archer, N., Duesing, K., Hannan, G., and Keast, R. (2016) Mechanism of fat taste perception: Association with diet and obesity. Prog Lipid Res 63, 41–49

2. Erlanson-Albertsson, C. (2010) Fat-Rich Food Palatability and Appetite Regulation. in Fat Detection: Taste, Texture, and Post Ingestive Effects (Montmayeur, J. P., and le Coutre, J. eds.), Boca Raton (FL). pp

3. Sullivan, E. L., Riper, K. M., Lockard, R., and Valleau, J. C. (2015) Maternal high-fat diet programming of the neuroendocrine system and behavior. Horm Behav 76, 153–161

4. Rivera, H. M., Christiansen, K. J., and Sullivan, E. L. (2015) The role of maternal obesity in the risk of neuropsychiatric disorders. Front Neurosci 9, 194

5. Karatayev, O., Lukatskaya, O., Moon, S. H., Guo, W. R., Chen, D., Algava, D., Abedi, S., and Leibowitz, S. F. (2015) Nicotine and ethanol co-use in Long-Evans rats: Stimulatory effects of perinatal exposure to a fat-rich diet. Alcohol 49, 479–489

6. Kang, S. S., Kurti, A., Fair, D. A., and Fryer, J. D. (2014) Dietary intervention rescues maternal obesity induced behavior deficits and neuroinflammation in offspring. J Neuroinflammation 11, 156

7. Hokfelt, T., Bartfai, T., and Bloom, F. (2003) Neuropeptides: opportunities for drug discovery. Lancet Neurol 2, 463–472

8. Barson, J. R., Morganstern, I., and Leibowitz, S. F. (2012) Neurobiology of consummatory behavior: mechanisms underlying overeating and drug use. ILAR J 53, 35–58

9. Barson, J. R., Morganstern, I., and Leibowitz, S. F. (2011) Similarities in hypothalamic and mesocorticolimbic circuits regulating the overconsumption of food and alcohol. Physiol Behav 104, 128–137

10. Adams, A. C., Clapham, J. C., Wynick, D., and Speakman, J. R. (2008) Feeding behaviour in galanin knockout mice supports a role of galanin in fat intake and preference. J Neuroendocrinol 20, 199–206

11. Hokfelt, T., Barde, S., Xu, Z. D., Kuteeva, E., Ruegg, J., Le Maitre, E., Risling, M., Kehr, J., Ihnatko, R., Theodorsson, E., Palkovits, M., Deakin, W., Bagdy, G., Juhasz, G., Prud’homme, H. J., Mechawar, N., Diaz-Heijtz, R., and Ogren, S. O. (2018) Neuropeptide and Small Transmitter Coexistence: Fundamental Studies and Relevance to Mental Illness. Front Neural Circuits 12, 106

12. Kormos, V., and Gaszner, B. (2013) Role of neuropeptides in anxiety, stress, and depression: from animals to humans. Neuropeptides 47, 401–419

13. Lin, E. J. (2012) Neuropeptides as therapeutic targets in anxiety disorders. Curr Pharm Des 18, 5709–5727

14. Madaan, V., and Wilson, D. R. (2009) Neuropeptides: relevance in treatment of depression and anxiety disorders. Drug News Perspect 22, 319–324

15. Karatayev, O., Baylan, J., and Leibowitz, S. F. (2009) Increased intake of ethanol and dietary fat in galanin overexpressing mice. Alcohol 43, 571–580

16. Karatayev, O., Baylan, J., Weed, V., Chang, S., Wynick, D., and Leibowitz, S. F. (2010) Galanin knockout mice show disturbances in ethanol consumption and expression of hypothalamic peptides that stimulate ethanol intake. Alcohol Clin Exp Res 34, 72–80

17. Lewis, M. J., Rada, P., Johnson, D. F., Avena, N. M., Leibowitz, S. F., and Hoebel, B. G. (2005) Galanin and alcohol dependence: neurobehavioral research. Neuropeptides 39, 317–321

18. Rada, P., Avena, N. M., Leibowitz, S. F., and Hoebel, B. G. (2004) Ethanol intake is increased by injection of galanin in the paraventricular nucleus and reduced by a galanin antagonist. Alcohol 33, 91–97

19. Davidson, S., Lear, M., Shanley, L., Hing, B., Baizan-Edge, A., Herwig, A., Quinn, J. P., Breen, G., McGuffin, P., Starkey, A., Barrett, P., and MacKenzie, A. (2011) Differential activity by polymorphic variants of a remote enhancer that supports galanin expression in the hypothalamus and amygdala: implications for obesity, depression and alcoholism. Neuropsychopharmacology 36, 2211–2221

20. McEwan, A. R., Davidson, C., Hay, E. A., Turnbull, Y., Erickson, J. C., Marini, P., Wilson, D., McIntosh, A. M., Adams, M. J., Murgatroyd, C., Barrett, P., Delibegovic, M., Clarke, T. K., and MacKenzie, A. (2020) CRISPR disruption and UK Biobank analysis of a highly conserved polymorphic enhancer suggests a role in male anxiety and ethanol intake. Molecular Psychiatry In Press

21. Murgatroyd, C., Patchev, A. V., Wu, Y., Micale, V., Bockmuhl, Y., Fischer, D., Holsboer, F., Wotjak, C. T., Almeida, O. F., and Spengler, D. (2009) Dynamic DNA methylation programs persistent adverse effects of early-life stress. Nat Neurosci 12, 1559–1566

22. Hay, E. A., Cowie, P., McEwan, A. R., Ross, R., Pertwee, R. G., and MacKenzie, A. (2020) Disease-associated polymorphisms within the conserved ECR1 enhancer differentially regulate the tissue-specific activity of the cannabinoid-1 receptor gene promoter; implications for cannabinoid pharmacogenetics. Hum Mutat 41, 291–298

23. Liu, C., Marioni, R. E., Hedman, A. K., Pfeiffer, L., Tsai, P. C., Reynolds, L. M., Just, A. C., Duan, Q., Boer, C. G., Tanaka, T., Elks, C. E., Aslibekyan, S., Brody, J. A., Kuhnel, B., Herder, C., Almli, L. M., Zhi, D., Wang, Y., Huan, T., Yao, C., Mendelson, M. M., Joehanes, R., Liang, L., Love, S. A., Guan, W., Shah, S., McRae, A. F., Kretschmer, A., Prokisch, H., Strauch, K., Peters, A., Visscher, P. M., Wray, N. R., Guo, X., Wiggins, K. L., Smith, A. K., Binder, E. B., Ressler, K. J., Irvin, M. R., Absher, D. M., Hernandez, D., Ferrucci, L., Bandinelli, S., Lohman, K., Ding, J., Trevisi, L., Gustafsson, S., Sandling, J. H., Stolk, L., Uitterlinden, A. G., Yet, I., Castillo-Fernandez, J. E., Spector, T. D., Schwartz, J. D., Vokonas, P., Lind, L., Li, Y., Fornage, M., Arnett, D. K., Wareham, N. J., Sotoodehnia, N., Ong, K. K., van Meurs, J. B. J., Conneely, K. N., Baccarelli, A. A., Deary, I. J., Bell, J. T., North, K. E., Liu, Y., Waldenberger, M., London, S. J., Ingelsson, E., and Levy, D. (2018) A DNA methylation biomarker of alcohol consumption. Mol Psychiatry 23, 422–433

24. Wynick, D., Small, C. J., Bacon, A., Holmes, F. E., Norman, M., Ormandy, C. J., Kilic, E., Kerr, N. C., Ghatei, M., Talamantes, F., Bloom, S. R., and Pachnis, V. (1998) Galanin regulates prolactin release and lactotroph proliferation. Proc Natl Acad Sci U S A 95, 12671–12676

25. Niehrs, C., and Calkhoven, C. F. (2020) Emerging Role of C/EBPbeta and Epigenetic DNA Methylation in Ageing. Trends Genet 36, 71–80

26. Angeloni, A., and Bogdanovic, O. (2019) Enhancer DNA methylation: implications for gene regulation. Essays Biochem 63, 707–715

27. Avrahami, D., and Kaestner, K. H. (2019) The dynamic methylome of islets in health and disease. Mol Metab 27S, S25–S32

28. Keleher, M. R., Zaidi, R., Hicks, L., Shah, S., Xing, X., Li, D., Wang, T., and Cheverud, J. M. (2018) A high-fat diet alters genome-wide DNA methylation and gene expression in SM/J mice. BMC Genomics 19, 888

29. Stadler, M. B., Murr, R., Burger, L., Ivanek, R., Lienert, F., Scholer, A., Wirbelauer, C., Oakeley, E. J., Gaidatzis, D., Tiwari, V. K., and Schubeler, D. (2011) DNA-binding factors shape the mouse methylome at distal regulatory regions. Nature 480, 490–495

30. Jones, P. L., Veenstra, G. J., Wade, P. A., Vermaak, D., Kass, S. U., Landsberger, N., Strouboulis, J., and Wolffe, A. P. (1998) Methylated DNA and MeCP2 recruit histone deacetylase to repress transcription. Nat Genet 19, 187–191

31. Nan, X., Ng, H. H., Johnson, C. A., Laherty, C. D., Turner, B. M., Eisenman, R. N., and Bird, A. (1998) Transcriptional repression by the methyl-CpG-binding protein MeCP2 involves a histone deacetylase complex. Nature 393, 386–389

32. Thurman, R. E., Rynes, E., Humbert, R., Vierstra, J., Maurano, M. T., Haugen, E., Sheffield, N. C., Stergachis, A. B., Wang, H., Vernot, B., Garg, K., John, S., Sandstrom, R., Bates, D., Boatman, L., Canfield, T. K., Diegel, M., Dunn, D., Ebersol, A. K., Frum, T., Giste, E., Johnson, A. K., Johnson, E. M., Kutyavin, T., Lajoie, B., Lee, B. K., Lee, K., London, D., Lotakis, D., Neph, S., Neri, F., Nguyen, E. D., Qu, H., Reynolds, A. P., Roach, V., Safi, A., Sanchez, M. E., Sanyal, A., Shafer, A., Simon, J. M., Song, L., Vong, S., Weaver, M., Yan, Y., Zhang, Z., Zhang, Z., Lenhard, B., Tewari, M., Dorschner, M. O., Hansen, R. S., Navas, P. A., Stamatoyannopoulos, G., Iyer, V. R., Lieb, J. D., Sunyaev, S. R., Akey, J. M., Sabo, P. J., Kaul, R., Furey, T. S., Dekker, J., Crawford, G. E., and Stamatoyannopoulos, J. A. (2012) The accessible chromatin landscape of the human genome. Nature 489, 75–82

33. Yin, Y., Morgunova, E., Jolma, A., Kaasinen, E., Sahu, B., Khund-Sayeed, S., Das, P. K., Kivioja, T., Dave, K., Zhong, F., Nitta, K. R., Taipale, M., Popov, A., Ginno, P. A., Domcke, S., Yan, J., Schubeler, D., Vinson, C., and Taipale, J. (2017) Impact of cytosine methylation on DNA binding specificities of human transcription factors. Science 356

34. Zhu, T., Brown, A. P., and Ji, H. (2020) The Emerging Role of Ten-Eleven Translocation 1 in Epigenetic Responses to Environmental Exposures. Epigenet Insights 13, 2516865720910155

35. Spruijt, C. G., Gnerlich, F., Smits, A. H., Pfaffeneder, T., Jansen, P. W., Bauer, C., Munzel, M., Wagner, M., Muller, M., Khan, F., Eberl, H. C., Mensinga, A., Brinkman, A. B., Lephikov, K., Muller, U., Walter, J., Boelens, R., van Ingen, H., Leonhardt, H., Carell, T., and Vermeulen, M. (2013) Dynamic readers for 5-(hydroxy)methylcytosine and its oxidized derivatives. Cell 152, 1146–1159

36. Stroud, H., Feng, S., Morey Kinney, S., Pradhan, S., and Jacobsen, S. E. (2011) 5-Hydroxymethylcytosine is associated with enhancers and gene bodies in human embryonic stem cells. Genome Biol 12, R54

37. Hon, G. C., Rajagopal, N., Shen, Y., McCleary, D. F., Yue, F., Dang, M. D., and Ren, B. (2013) Epigenetic memory at embryonic enhancers identified in DNA methylation maps from adult mouse tissues. Nat Genet 45, 1198–1206

38. Hashimoto, H., Olanrewaju, Y. O., Zheng, Y., Wilson, G. G., Zhang, X., and Cheng, X. (2014) Wilms tumor protein recognizes 5-carboxylcytosine within a specific DNA sequence. Genes Dev 28, 2304–2313

39. Sun, Z., Xu, X., He, J., Murray, A., Sun, M. A., Wei, X., Wang, X., McCoig, E., Xie, E., Jiang, X., Li, L., Zhu, J., Chen, J., Morozov, A., Pickrell, A. M., Theus, M. H., and Xie, H. (2019) EGR1 recruits TET1 to shape the brain methylome during development and upon neuronal activity. Nat Commun 10, 3892

